# Targeting the mTORC2 signaling complex in B cell malignancies

**DOI:** 10.1101/564500

**Authors:** Wei Liao, Gwen Jordaan, Angelica Benavides-Serrato, Brent Holmes, Joseph Gera, Sanjai Sharma

## Abstract

Hyperactive PI3 kinase-Akt (PI3K-Akt) signaling has an important role in cell growth and resistance to apoptosis in B cell malignancies. Inhibition of this pathway by blocking PI3K activity, and or inhibiting mTORC1/2 signaling complexes is an active area of research in B cell leukemia/lymphoma such as chronic lymphocytic leukemia (CLL) and mantle cell lymphoma (MCL). With a tissue-scan array, the expression of Rictor is a component of the mTORC2 complex was determined by quantitative PCR in a number of B cell malignancies. Rictor was found to be over-expressed in CLL and MCL cells as compared to normal B cells with no over-expression in Hodgkins and non-Hodgkins lymphomas. Inactivation of Rictor was performed by shRNA in two Mantle cell lines and these stable Rictor knockdown cell lines demonstrated a slower growth of cells as compared to scrambled shRNA control. In addition, there was a decrease of mTORC2 signaling and B cell receptor (BCR) cross-linking mediated Akt (Ser473) and NDRG1 (Thr 346) phosphorylation. To specifically disrupt the mTORC2 signaling complex and target Rictor overexpression, previously identified inhibitors that block Rictor and MTOR interaction in a yeast two-hybrid system were analyzed. Treatment of primary CLL specimens with these inhibitors followed by immunoprecipitation experiments confirmed the disruption of the mTORC2 complex. These inhibitors also induced apoptosis in CLL specimens and were more effective than rapamycin, an MTOR inhibitor and pp242, an mTORC1 and 2 inhibitors, at equimolar concentrations. Treatment of CLL specimens with the lead inhibitor, compound#6, resulted in inhibition of p-Akt, p-GSK 3 beta, p-PKC alpha, p-Foxo1, and p-Foxo3, with minimal effect on the phosphorylation of an mTORC1 target gene, S6 kinase. In comparison with Idelalisib (CAL-101), a clinically approved PI3Kinase p110 delta inhibitor in CLL, comp#6 is more effective in inducing apoptosis in primary CLL specimens at equimolar concentrations (mean 51.2, SD 21.7 as compared to mean 26.9, SD 17.2). The data support the effectiveness of these novel inhibitors that specifically disrupt the mTORC2 complex in primary CLL specimens.

## Introduction

CLL is a B cell malignancy characterized by a steady accumulation of leukemic B cells as these cells exhibit resistance to apoptosis. B-cell receptor (BCR) signaling is the major signaling pathway and defines clinical, biologic and prognostic characteristics of this disease [1,2]. Signaling via BCR leads to cell proliferation, survival, and resistance to apoptosis [1,2]. The activation or cross-linking of BCR results in the formation of a signaling complex (signalosome) that includes Lyn, Syk, BTK (Bruton tyrosine kinase), and Zap-70 among other components. This complex in turn activates a number of downstream pathways such as PI3Kinase/Akt/mTOR (PI3K/Akt/mTOR), PKC (protein kinase C), NFKB (nuclear factor kappa B) and ERK (extracellular signal-regulated kinase) that drive cell proliferation, resistance to apoptosis, cell motility, migration, etc. [3–7]. Besides BCR initiated signaling, additional signaling by microenvironment also has profound effects on the growth and resistance to apoptosis of these cells. This includes signaling via chemokines (CCL3, CCL4, and CXCL13), cytokines (IL4), tumor necrosis factor receptor, CD40 ligand, and other stromal factors that activate growth and survival pathways in these cells [8].

PI3Kinase is a heterodimer of a regulatory subunit, p85 and a p110 catalytic subunit that generate the second messenger phosphatidylinositol-3,4,5-triphosphate. This occurs at lipid rafts where Akt, a serine-threonine kinase is recruited and phosphorylated by phosphoinositide-dependent kinase-1 (PDK-1). Akt promotes cell proliferation, survival and migration and its complete activation requires phosphorylation by PDK-1 and the mTORC2 complex. mTOR is a serine-threonine kinase that exists in two complexes, mTORC1, and mTORC2 which regulate distinct cellular processes. mTORC1 is composed of mTOR, raptor, mLST8, and PRAS40, and upon activation inhibits 4E-BP1 (eIF4E inhibitory binding protein) and promotes cap-dependent translation and protein synthesis while the mTORC2 complex includes mTOR, Rictor, mLST8, mSin2 and phosphorylates AKT, PKC alpha, and SGK1 [9–11]. The activation of PI3K/Akt/mTOR overall results in upregulation of anti-apoptotic genes such as Mcl1 and XIAP that promote survival of leukemic cells [7,12,13]. In addition, there is phosphorylation and inactivation of BAD, a pro-apoptotic gene of the Bcl2 family, Forkhead family of transcription factors and GSK3β, a negative regulator of cyclins and MYC [14,15,16].

With its role in growth regulation, apoptosis, and survival, a number of strategies to inhibit PI3 kinase/Akt at multiple levels have been attempted. CAL-101 (Idelalisib) is a PI3K p110, delta isoform inhibitor that inhibits Akt activation along with inhibition of important CLL-microenvironment interactions and is in clinical use to treat CLL [17]. Besides PI3Kinase inhibitors, there are specific inhibitors that block mTOR function including Rapamycin and its analogs RAD001 (everolimus) and CCI-779 (temsirolimus) which are inhibitors primarily of mTORC1 complex and act via the immunophilin FK506-binding protein (FKBP12)[18]. In addition there are inhibitors of mTORC1/2 and dual inhibitors of both mTORC1 and 2 [19–21]. The Akt/PI3K pathway is also active in another B cell malignancy, Mantle cell lymphoma [22,23]. The therapeutic efficacy of the currently available mTORC1 inhibitors in CLL and Mantle cell lymphoma is low due to a negative feedback loop by which mTORC1 inhibition leads to AKT activation through upregulation of receptor tyrosine kinases (RTKs, or substrates) and also Erk pathway activation via IRS-1 [24–26]. On the contrary, mTORC2 directly activates AKT and therefore mTORC2-specific inhibitors potentially will be more effective anti-cancer drugs. Such compounds would block Akt phosphorylation and not disrupt the mTORC1-dependent negative feedback loops, which promote drug resistance to mTORC1 inhibitors.

The aim of this study was to investigate the effectiveness of previously identified chemical compounds that block Rictor-MTOR protein-protein interaction in B cell malignancies. These compounds were identified by a yeast two-hybrid screening and the methodology has reported by Benavides-Serrato et al [27]. We chose the B cell malignancies to analyze these compounds as the PI3K/Akt/mTOR pathway is highly active in these malignancies. All CLL experiments were performed on primary leukemic cells and the data demonstrates that these compounds inhibit this pathway and growth of this B cell leukemia.

## Materials and Methods

### Primary CLL specimens and cell lines

Peripheral blood samples were obtained from CLL patients at the Los Angeles VA hospital after informed written consent and institutional approval. Leukemic cells were isolated by ficoll density gradient using Ficoll-Paque™ Premium (GE Healthcare) and stored in liquid nitrogen. CLL cells were cultured in RPMI 1640 (Mediatech) containing 10% FCS, 1 mM sodium pyruvate, 2 mM L-glutamine and penicillin-streptomycin solution. PBMCs from normal donors were isolated using a ficoll density gradient, and CD19+ B-cells were isolated by negative selection using the Dynabeads^®^ Untouched™ Human B Cells kit (Invitrogen). Mantle cell lines (Z-138, Rec-1, and Maver-1) were purchased from ATCC.

### Plasmids, transfections, and reagents

The shRNA/pLOK.1 targeting rictor (plasmid #1853, Addgene) and the control scrambled shRNA (plasmid #1864, Addgene) was provided by Dr. Alan Lichtenstein (UCLA West Los Angeles and VA Medical Center, Los Angeles, CA). Mantle cell lines Z138 and Rec-1 were infected with lentiviral vectors with MOIs of 5, selected with 0.5 μg/ml puromycin and stable pools of cells were obtained. Rapamycin was purchased from Calbiochem, pp242, and Idelalisib (CAL101 or Zydelig, Gilead Sciences) from Selleckchem, both inhibitors were dissolved in DMSO. Chemical compounds blocking the protein-protein interaction between Rictor and mTOR (mTORC2 compounds) 4,6 and 7 were isolated via a yeast two-hybrid drug screen and provided by Dr. Joseph Gera (UCLA West Los Angeles and VA Medical Center, Los Angeles, CA, (manuscript submitted). All antibodies, except p70-S6 kinase (Abcam) and actin (Santa Cruz Biotechnologies), were purchased from Cell Signaling.

### Real-Time PCR and qPCR Array Analysis

RNA from CLL samples and B-cells was isolated by the RNAeasy^®^ Mini kit by Qiagen. cDNA was synthesized using oligo-dT, dNTP and SuperScript II Reverse Transcriptase from Invitrogen. The TissueScan™ Tissue qPCR Array containing cDNAs for human lymphoma I-II was purchased from Origene. Relative expression of Rictor was determined using inventoried Taqman probes and PCR master mix from Applied Biosystems. Real-time PCR expression of actin was used as an endogenous control, and relative expression was calculated as described by Pfaffl [28].

### Cell Survival and Apoptosis Assays

Z138 and Rec-1 cell lines with Rictor knockdown or scrambled shRNA control were seeded at 5×10^4^ cells and assayed for cell growth over 8 days. Live cells were enumerated in triplicate by trypan blue negative staining. Apoptosis in cell lines and CLL cells was measured by Annexin staining and flow cytometry using the FITC Annexin V Apoptosis Detection Kit 1 (BD Pharmingen). Percentage apoptosis represents the % of apoptotic cells above the % apoptosis in control cells. Apoptosis of CLL specimens was also tested in two different culture condition, with media, and with a stromal cell line (HS-5, ref 29) co-culture to mimic microenvironment support.

### Immunoprecipitation and Immunoblotting

Cells were lysed in CST (Cell Signaling Technology) lysis buffer supplemented with protease inhibitor cocktail (Pierce) and quantified (BioRad DC protein assay). 30 μg of lysate was loaded to SDS-polyacrylamide gels. Blots were probed with primary antibodies followed by HRP-conjugated secondary antibodies and signal detected by GE-Amersham chemiluminescent kit and Fuji- LAS imager. Immunoprecipitation of the mTORC2 complex was performed by lysing cells in the CHAPS buffer (CST) and immunoprecipitating complexes by an anti-Rictor antibody. The precipitate was immunoblotted for mTOR. Lysates were also analyzed by mTOR and Rictor antibodies on western blots. Immunoprecipitation studies were also designed to assess changes in P-AKT signals with mTORC2 compounds by immunoprecipitating total Akt followed by western blots with a P-Akt antibody.

## Results

### Rictor expression in lymphoma and CLL specimens

mTORC2 complex regulates cell growth and survival signals and in contrast to mTORC1, it is nutrient-insensitive and has a role in cancer cell growth and resistance to apoptosis. Rictor is a component only of the mTORC2 complex along with MTOR, MLST8, PRR5, MAPKAP1, and DEPTOR and its over-expression can result in tumor formation and an increase in mTORC2 signaling [30–32]. To determine its expression in CLL and B-cell lymphoma specimens a quantitative real-time PCR analysis was performed. As depicted in Figure 1A, the relative Rictor RNA expression (actin control) is 0.2 to 2 fold higher in the majority of CLL specimens as compared to normal B cells. To determine expression in a number of additional B cell malignancies and primary lymphoma specimens a tissue scan array was analyzed. Rictor expression relative to normal lymph node cDNA was higher in small lymphocytic leukemia (SLL), a CLL related B cell malignancy and Mantle cell lymphoma (MCL) specimens. Other lymphomas including Hodgkin’s, Follicular and diffuse large B cell lymphomas did not show Rictor over-expression in this assay. Of a number of B cell leukemia/lymphomas tested, CLL and Mantle cell lymphoma are thus two B leukemia/lymphomas cell with high Rictor expression and are also known for hyperactive PI3/Akt signaling pathway. On western blot analysis, the increased expression of Rictor as compared to peripheral blood normal B cells is more clearly evident as compared to the RNA expression (figure 1C).

**Figure 1:**
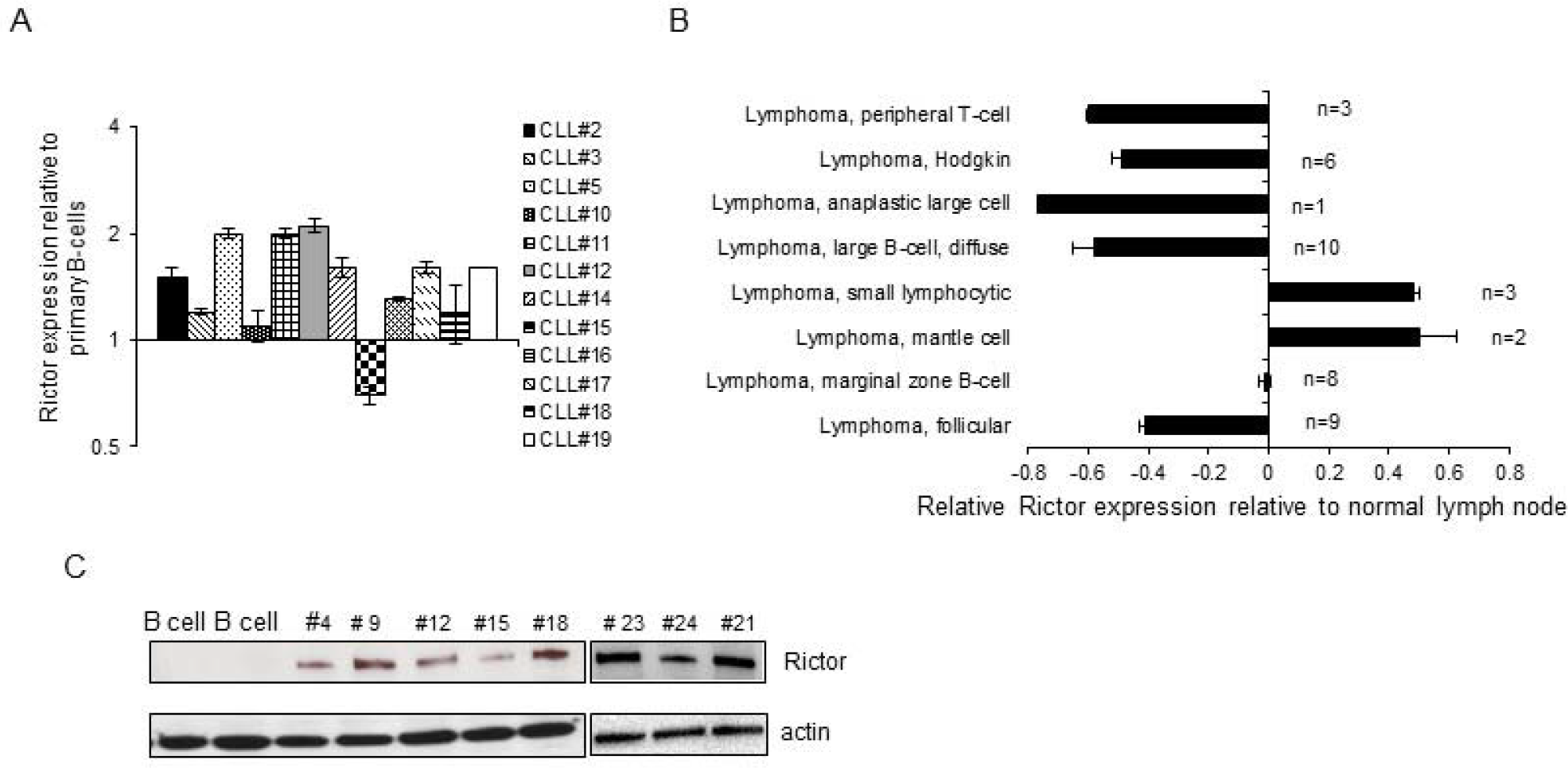
Rictor expression in B-cell lymphoproliferative disorders. A. Real-time PCR data. Twelve primary CLL specimens were analyzed for Rictor expression by real-time PCR using Taqman probes and expression determined relative to CD19+ primary human peripheral blood B cells. (actin control). B. A tissuescan PCR array with cDNA from different B cell malignancies was analyzed by real-time PCR and the relative expression normalized to expression in normal lymph node cells. N denotes the number of specimens. C. Western blot analysis of Rictor expression in primary CLL specimens and magnetic bead isolated peripheral blood normal B cells (left two lanes). Rictor expression (192kd band) and actin control.

### Rictor knockdown in Mantle cell lines

To test the role of Rictor over-expression in B cells, a Rictor knockdown in Mantle cell lines (MCL) was performed as they have a high expression (Figure 1B). MCL cell lines are easy to transfect and can be activated by B-cell receptor cross-linking similar to primary CLL specimens. Four MCL lines were initially analyzed for Rictor expression by western blot and a strong Rictor band was detected (Figure 2A). Z138 and Rec-1 MCL lines were infected with lentiviral particles expressing Rictor shRNA or a scrambled control shRNA (Ctrl shRNA) and selected. Stably selected pools of Rictor knockdown cells and scrambled control cells were obtained. Rictor knockdown was confirmed by a lower Rictor expression by western blot analysis in the Rictor knockdown cells (Figure 2B). To analyze mTORC2 signaling in these Rictor knockdown cells, B-cell receptor was cross-linked by anti-IgM antibody followed by detection of phosphorylated AKT (Ser473) and phosphorylated NDRG1 (Thr 346). Both these proteins are phosphorylated by the mTORC2 complex and a change in the activity of this complex will be reflected by an alteration in the phosphorylation of target genes. The experiment was performed with and without cross-linking in the two established knockdown cell lines (figure 2C). Z138 cells have high baseline levels of P-AKT and P-NDRG1 signals that are diminished in Rictor knockdown cells. Also with cross-linking of the B-cell receptor, the phosphorylation of AKT (Ser473) and NDRG1 (Thr 346) is decreased in Rictor knockdown cell lines as compared to the Ctrl scrambled shRNA lines. In Rec-1 cells the p-AKT signal in non-anti-IgM treated cells is not observed but similar to Z138, there is a decrease in both AKT and NDRG1 phosphorylation with Rictor knockdown. To further analyze the effect of knockdown on growth characteristics of the cell, a growth curve assay was performed. In a cell count assay over an eight-day period, the Rictor knockdown cell lines (Z138 and Rec-1) grow slower as compared to the Ctrl shRNA cell lines (figure 2D). Overall the data shows that in the MCL model, Rictor is over-expressed and its expression is required for cell growth and the functional activity of the mTORC2 complex which is diminished in cells with lower Rictor expression.

**Figure 2:**
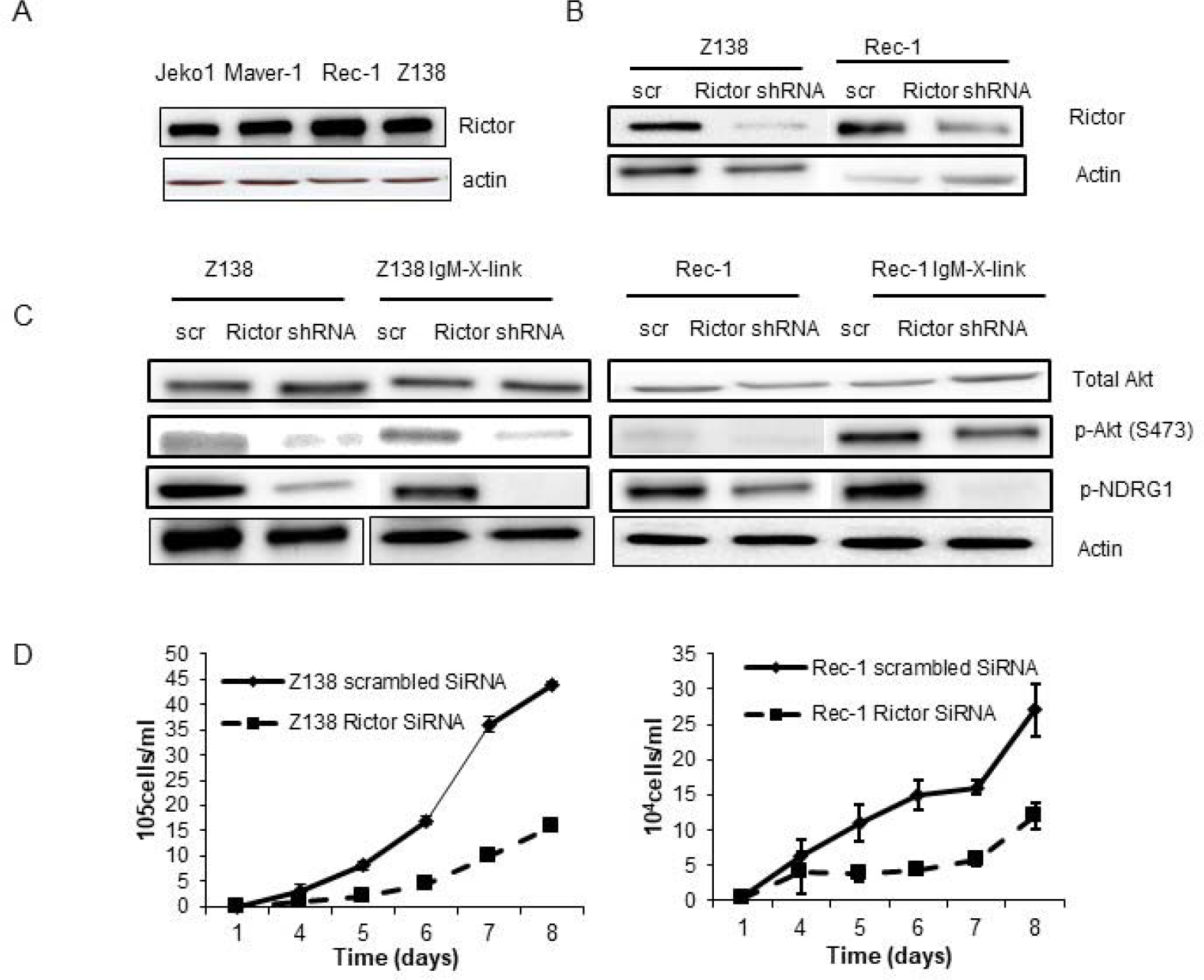
Rictor inactivation in Mantle cell lines. A. Rictor expression in Mantle cell lines, JeKo-1, Maver-1, Rec-1, and Z138. Cell lysates were analyzed for Rictor and actin expression by western blots. B and C, Z138 and Rec-1 mantle cell lines were infected with either a lentiviral vector expressing Rictor shRNA or a lentiviral vector expressing scrambled shRNA (control, Ctrl). Selected pools of cells were analyzed for Rictor expression by western blot analysis. C. Rictor shRNA or Ctrl selected cells were analyzed for mTORC2 signaling with IgM cross-linking. Lysates from cells with and without IgM X-linking (cross-linking with anti-IgM antibody at 10 μg/ml for 20 minutes) were analyzed for total Akt, phospho-Akt (Ser573) and phospho-NDRG1 (Thr346). D. The growth characteristics of Ctrl shRNA and Rictor shRNA expressing cells of Z138 and Rec-1 cell lines were analyzed by counting cells over a period 8 days.

### Apoptosis with PI3Kinase/AKT pathway inhibitors

As Rictor expression is higher in B cell lymphomas and leukemia and its downregulation inhibits mTORC2 signaling, we next tested the effect of compounds that inhibit MTOR-Rictor interaction. These compounds were isolated by a yeast two-hybrid screen and inhibit MTOR-Rictor binding. As Rictor is present only in the mTORC2, the compounds potentially disrupt the complex and thereby selectively inhibit mTORC2 function [27]. These compounds labeled, #4,6 and 7 were tested for apoptosis induction along with other known inhibitors of this pathway, namely, Rapamycin, an MTOR inhibitor and pp242 a dual MTORC1/2 inhibitor. For comparison, all inhibitors were analyzed at 1μM concentration and apoptosis was measured by flow cytometry-based annexin assay 48 hours after treatment. Data from four primary CLL specimens is shown in figure 3A, with compound #6 being the most effective in inducing apoptosis. Rapamycin and pp242 did not induce significant apoptosis in these cells. Based on this screening assay, compound #6 was selected for further analysis and its chemical structure is shown in figure 3B.

**Figure 3:**
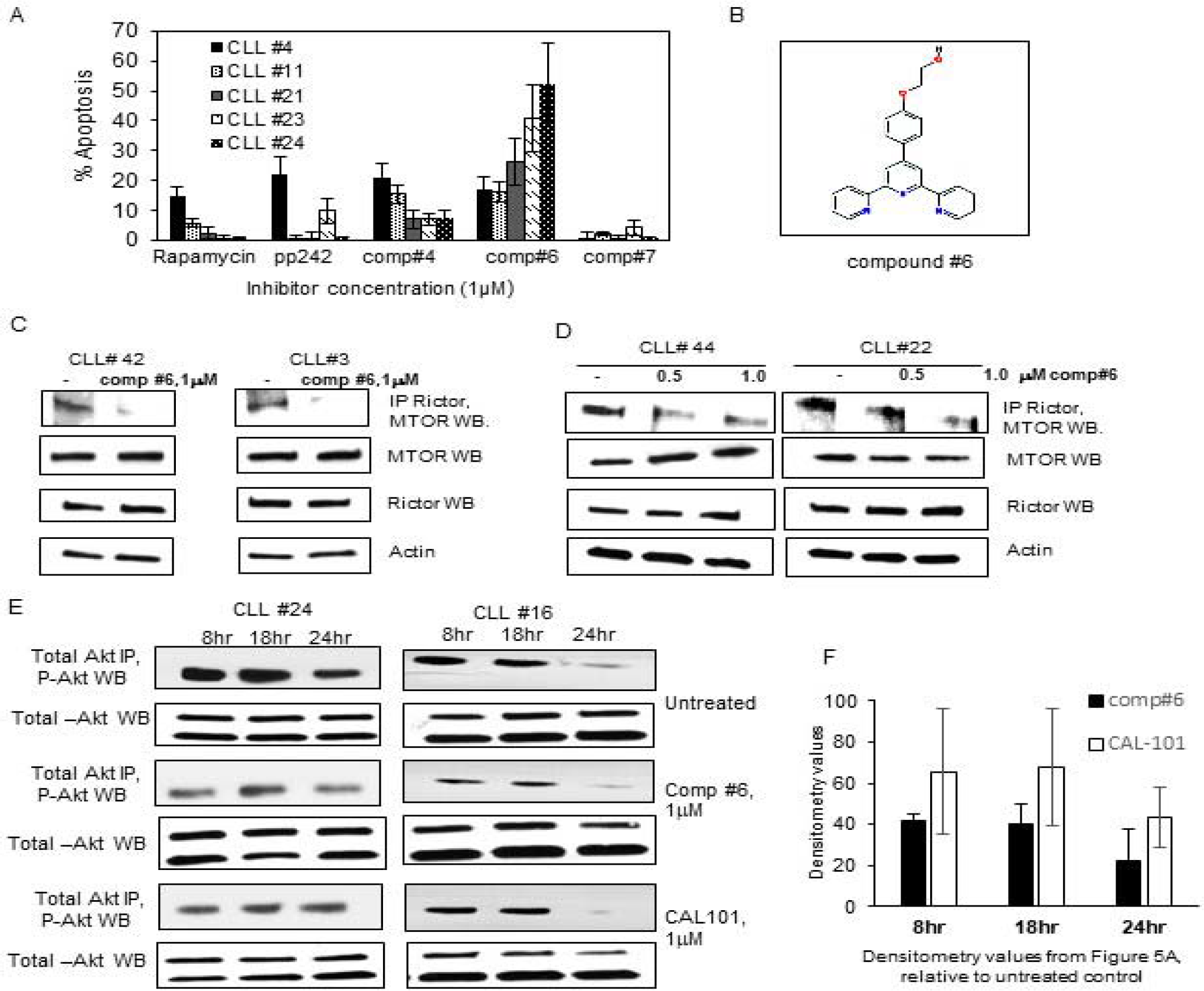
A. Induction of apoptosis by Rapamycin, pp242, and Rictor-MTOR inhibitors, compound #4, #6 and #7. Five primary CLL specimens were treated with different inhibitors at 1 μM concentration for 48 hours. Cells were analyzed for apoptosis using the Annexin flow assay and the data shown is apoptosis above background. Experiment repeated twice with similar results. B. Chemical structure of compound#6. C. Disruption of the mTORC2 complex with compound#6. Two primary CLL specimens were treated with compound#6 for 12 hours and lysates in CHAPS buffer were immunoprecipitated with the anti-Rictor antibody. The immunoprecipitate was loaded on an SDS gel and analyzed for MTOR expression by western blot analysis. In cells treated with compound#6, MTOR signal is much weaker as it cannot be immunoprecipitated with Rictor antibody. Western blots for MTOR, Rictor, and actin with identical lysates show equal amounts of these proteins. D. Two CLL specimens were treated with two different concentrations of compound#6 (0.5 and 1.0 μM) and analyzed by co-immunoprecipitation. MTORC2 complexes were immunoprecipitated with mTORC2 antibody followed by western blot with Rictor antibody. There is a dose-dependent decrease in the Rictor signal when cells are treated with compound#6. E. Two primary CLL specimens were treated with compound#6 at 1μM concentration for different time points. Total AKT was immunoprecipitated by an antibody the immunoprecipitate was analyzed by western blot analysis with P-AKT (Ser473). The experiment was also done identically with CAL-101 (1μM concentration). As compared to the untreated immunoprecipitates, compound#6 and CAL-101 specimens show a decrease in P-AKT signals. F. Densitometry representation of data in Figure 3E. Band intensity relative to the untreated control for each time point was given a value of 100. The bar diagram shows inhibition of phospho-Akt by compound#6 and CAL-101 at all the time points tested with comparatively higher inhibition by comp #6 at equimolar concentration.

To determine its mechanism of action, two primary CLL specimens were treated with compound#6 for 12 hours and the mTORC2 complexes were immunoprecipitated by a Rictor antibody. The immunoprecipitate was then analyzed by western blot for MTOR expression. Figure 3C, top panel, shows a decrease in the intensity of mTOR signal with compound#6 treatment in the two CLL specimens indicating a lack of physical association between Rictor and MTOR in the lysates. Lower panels in figure 3C are western blots for mTOR, Rictor using the same lysate with no change in overall expression of Rictor and mTOR. In two additional CLL specimens, the experiment was repeated with two different concentrations of compound#6 (Figure 3D) and as in Figure 3C and there was a decrease in mTOR western blot signal when mTORC2 complexes were immunoprecipitated with Rictor antibody in compound#6 treated CLL specimens. These co-immunoprecipitation experiments indicate that the compound#6 is able to decrease the association of Rictor and mTOR in CLL leukemic cells.

Compounds that inhibit the MTOR and Rictor interaction are expected to block activation of mTORC2 complex target genes including Akt. The phospho-AKT signal is weak in CLL specimens and not consistently observed in non-stimulated cells, therefore, experiment was modified to include an immunoprecipitation step with a total AKT antibody followed by a phospho-AKT (Ser473) western blot. Figure 3E shows western blot analysis of two CLL specimens that were treated with comp #6 and Idelalisib (PI3Kinase P110 delta inhibitor, 1μM concentration) for 8, 18 and 24 hours. An identical amount of lysates were immunoprecipitated by total AKT antibody (Figure 3E) and then analyzed for phospho-AKT (Ser473) by western blots. The western blot shows a decrease in the phospho-Akt (Ser473) signal with comp#6 and CAL-101. With no activation of B-cell receptor signaling, it is clearly evident that there is a baseline phosphorylation of AKT in CLL specimens that is clearly inhibited by blocking the PI3kinase/Akt pathway. Primary CLL specimens in culture demonstrate a decrease in phospho-AKT signal over time as the viability decreases, however, this signal is further decreased with compound#6 and Idelalisib. Figure 3F is a bar diagram with average densitometry values from figure 3E western blots. There is a 40% inhibition in the phospho-AKT signal relative to the untreated control at 8 and 18 hours of compound#6 treatment, with a further decline at the 24-hour time point. Idelalisib also decreased the phospho-AKT signal in unstimulated CLL specimens as expected, however, the percentage decrease is comparatively less as compared to compound#6. This series of western blots demonstrate that compound#6 blocks the Rictor-MTOR physical interaction in primary CLL specimens and inhibits Akt phosphorylation. Comparison of the immunoprecipitation assays performed at equimolar concentrations and identical time points, inhibition by Idelalisib is not as effective as compound#6. Some variation between specimens is expected as not all CLL specimens have high and active levels of the Idelalisib target, the PI3Kinase p110 delta while it is possible that an inhibitor blocking Rictor and MTOR interaction, two ubiquitous proteins will be consistently more effective in inhibiting this pathway.

### The activity of MTOR-Rictor inhibitors on primary CLL specimens

To characterize the signaling pathways affected by compound#6, two CLL leukemic specimens were treated with two concentrations of compound#6 for 12 hours and then B-cell receptor was activated by anti-IgM antibody for activation. Figure 4A shows western blots of two CLL specimens with and without BCR activation (IgM cross-linking) and with the comp#6 treatment at two concentrations. P-AKT signal is observed upon BCR activation as expected and in addition, BCR activation also phosphorylates a number of mTORC2 target genes including, NDRG1, GSK-3β (Ser9), PKC α, and FOXO3A (Ser253). Pre-treatment of CLL specimens with comp#6 inhibits this pathway activation in a dose-dependent manner as treated cells demonstrate decreased phosphorylation of mTORC2 target genes. Inhibition of phosphorylation of FOXO3A was observed in only one CLL specimen. In addition, two CLL specimens were treated for different time points (2,5,8,12, and 18 hours) at 1μM concentration of compound#6 and at each time point specimens were cross-linked with anti-IgM antibody for B-cell receptor activation (Figure 4B). This figure shows an effect of compound#6 on AKT pathway target genes in CLL specimens. Phospho-AKT downregulation is expected with mTORC2 inhibition and this was observed between the 5 to 12 hour time points in the two CLL specimens. The variability of phospho-AKT inhibition is likely due to the fact that BCR activation stimulates a number of pathways and it is possible that the S473 residue is phosphorylated by other kinases as well, and additionally these substrates are acted upon by a number of effectors of BCR activation. Similarly, there was inhibition of GSK-3β(Ser9), PKC-alpha and FOXO1 (Thr 24) with 5-8 hours of treatment. The phosphorylation of S6 kinase, a mTORC1 target in these compound#6 treated CLL specimens remains unchanged at different time points and indicates that this compound does not have an effect on mTORC1 signaling This is significant as inhibition of mTORC1 signaling results in activation of feedback loops that interfere with apoptosis and cell death.

**Figure 4:**
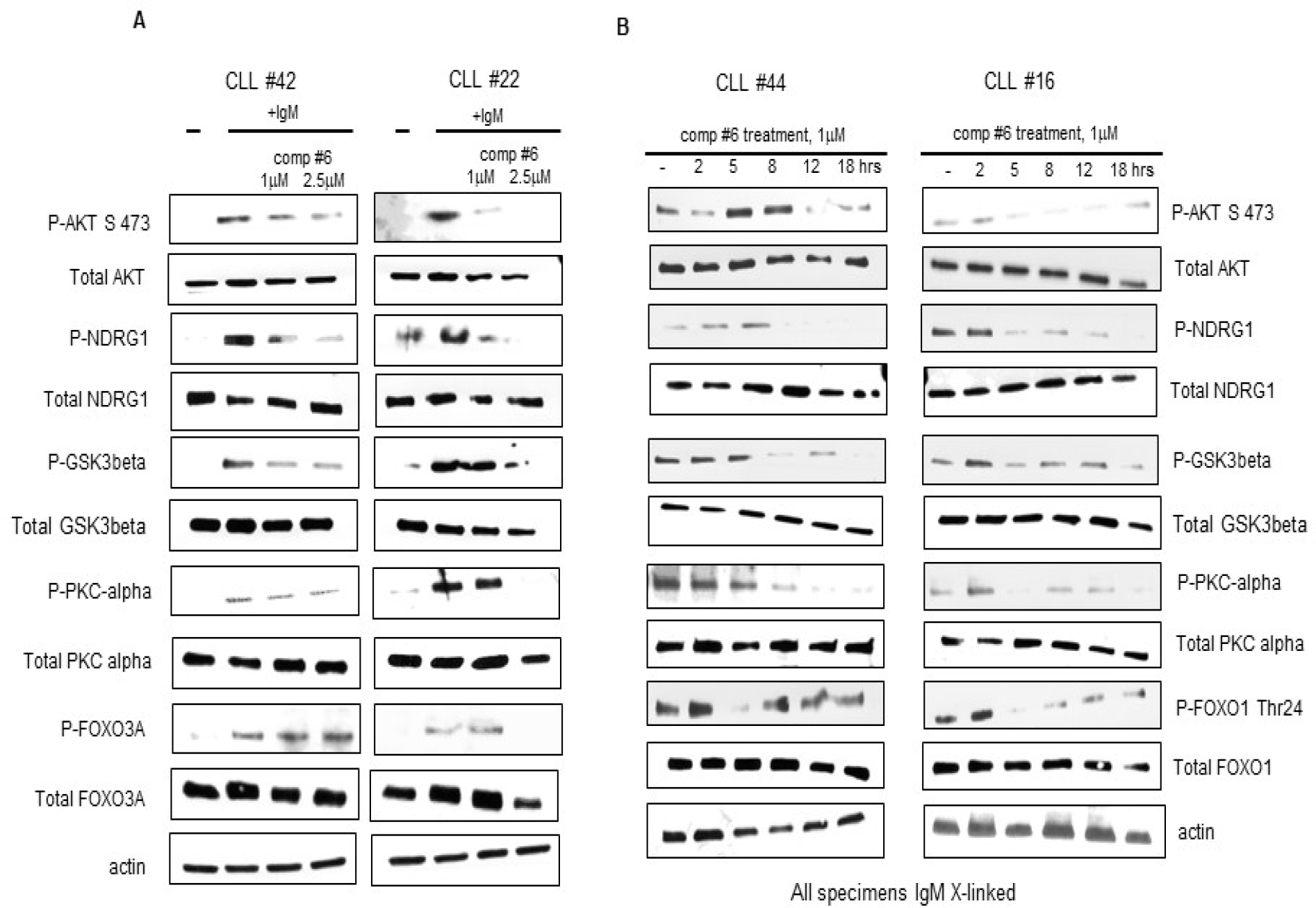
A. Two CLL specimens were treated with two concentrations of compound#6 for 12 hours and then cross-linked with IgM for BCR activation. The lanes are untreated CLL cells, un-treated and IgM activated cells, compound#6 treated and then IgM activated cells as indicated. Compound#6 inhibits the activation of a number of mTORC2 downstream signaling intermediates. B. Two CLL specimens were treated with compound#6 for different time points and then cross-linked (including the untreated cells, first lane) with IgM antibody as described. Lysates were analyzed by western blot analysis for expression of Total Akt, P-Akt (S473), P-NDRG1 (Thr346), P-S6 kinase (Thr389), P-GSK3β (Ser9), P-PKCα(Tyr658), P-Foxo1 (Thr24) and actin.

### MTORC2 inhibition and apoptosis

The AKT pathway is active in CLL specimens and there is a phospho-Akt signal in non-stimulated primary CLL specimens (Figure 3). Inhibition of this anti-apoptotic AKT signaling pathway alone is reportedly sufficient to induce apoptosis of CLL specimens [7]. To test this with compounds that block the Rictor-MTOR interaction, a number of CLL specimens were treated with a 1μM concentration of compound#6 and the lysates analyzed for Poly (ADP-ribose) polymerase (PARP) cleavage at 48 hours after treatment. Figure 5A is a western blot analysis of CLL specimens showing PARP cleavage by caspases that results in an increased intensity of the smaller (89kD) band with compound#6 treatment. To further study the activity of the compound#6, a number of CLL specimens were treated with this inhibitor and Idelalisib for a comparative analysis. In figure 5B, a table shows the data with % apoptosis induced by these inhibitors on 11 primary CLL specimens. Both inhibitors were used at 1μM concentration for 48 hours, and apoptosis detected by flow cytometry annexin assay. For all the CLL specimens tested, mean % apoptosis with compound#6 was 46%±22 (mean ± SD) and with Idelalisib was 26.9 ±17.2. Compound#6 was nearly twice as effective as Idelalisib in inducing apoptosis at identical concentration.

**Figure 5:**
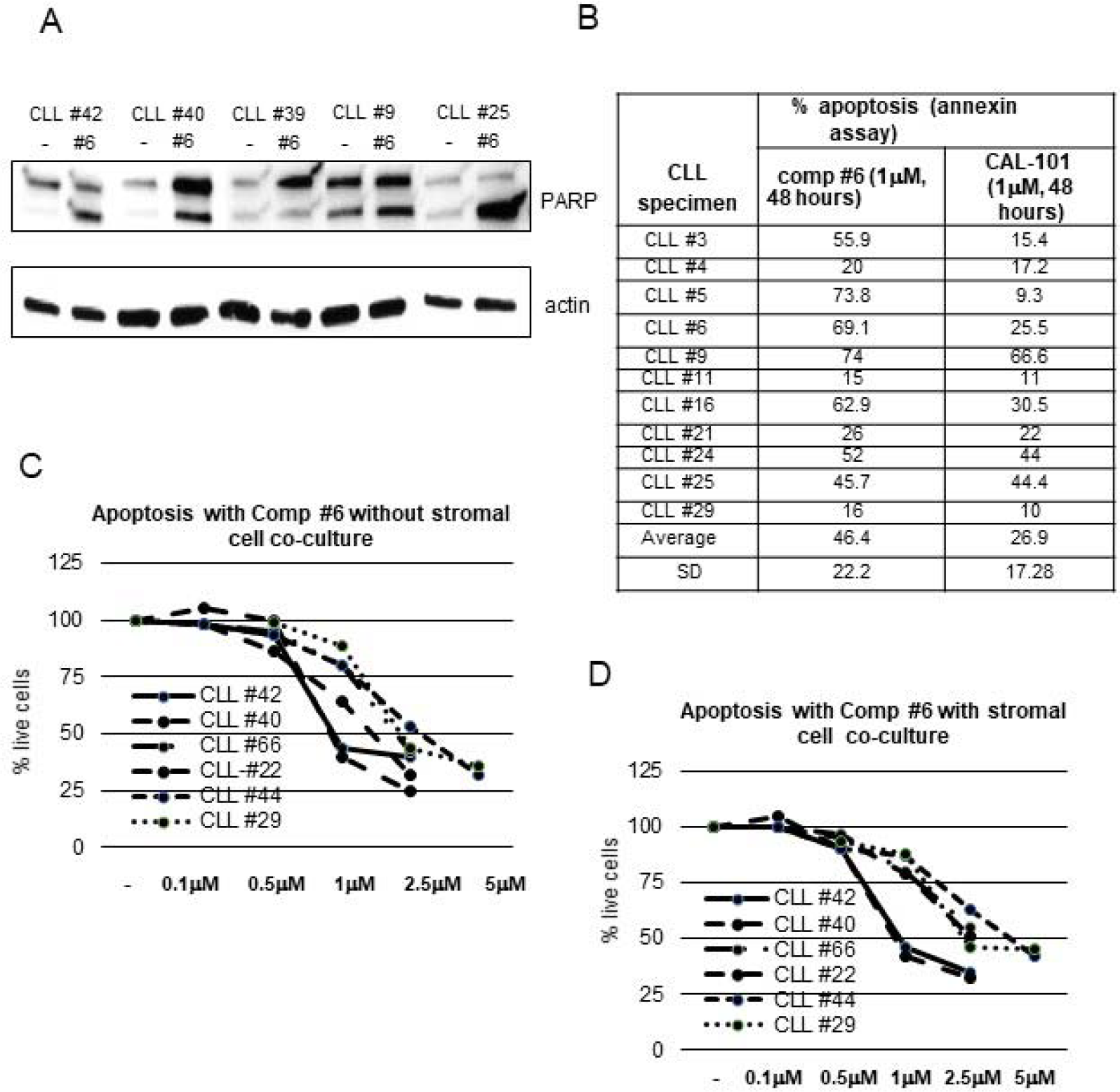
A. Western blot analysis of primary CLL specimens treated with 1 μM compound#6 for 48 hours and then analyzed for PARP cleavage by western blot analysis. The arrows indicate the upper (full-length) and lower (cleaved) PARP band that increases in intensity with apoptosis. B. Table with % apoptosis analyzed by an annexin flow-cytometry based assay. Primary CLL specimens were treated with compound#6 or CAL-101 at 1 μm concentration and analyzed at 48 hours after treatment. C. D. Six CLL specimens were treated with compound#6 at different concentrations without stromal cell co-culture (C) and with stromal cells (HS-5 cells, panel D). Apoptosis was analyzed at 48 hours by Annexin flow cytometry assay. The line diagrams indicate the % of live cells at various compound#6 concentrations.

A more clinically meaningful way to measure the effectiveness of a drug in CLL is to determine the degree of apoptosis in leukemic cells when they are cultured with stromal cells. These stromal cells provide a supportive microenvironment to these primary leukemic cells which is similar to the in vivo microenvironment. Previous studies have shown that inhibition of PI3/AKT pathway by Idelalisib also suppresses microenvironment signaling in CLL in addition to inducing apoptosis [17]. To determine whether there was a similar effect of compound#6, apoptosis experiments were performed on primary CLL specimens with and without co-culture with HS-5 stromal cells. Figure 5C, D shows data from apoptosis experiments on two CLL specimens with different concentrations of compound#6. In these experiments, there was a 10-20% higher viability of untreated cells when they were cultured with stromal cells as compared to growth media alone. Figure 5C, D indicates that compound#6 was able to induce apoptosis in these leukemic cells in a dose-dependent manner both with or without stromal cell co-culture. Induction of apoptosis in CLL leukemic cells co-cultured with stromal cells indicates that compound#6 is able to overcome PI3K/Akt signaling mediated via microenvironment and effectively counteract the protective effect of the microenvironment.

### Discussion

Rictor over-expression has been observed in a number of tumor model systems and correlates with a higher activity of the mTORC2 signaling complex and oncogenic properties of tumor cells [30–32]. This study identified that Rictor, a component of the mTORC2 complex is over-expressed in primary CLL specimens and mantle cell lymphoma [33–35]. In addition, knockdown of Rictor inhibits the growth of cells and downregulates mTORC2 signaling implying a critical role of this protein in the functioning of the mTORC2 complex. To specifically target Rictor and the mTORC2 complex, a number of inhibitors that prevent Rictor-MTOR interaction in yeast were tested on CLL specimens and were able to induce apoptosis in primary CLL specimens. The inhibitor were more effective than a mTOR inhibitor, dual mTORC1,2 inhibitor and a clinically approved PI3 kinase inhibitor, Cal-101, Idelalisib.

These small molecule inhibitors were isolated using a yeast two-hybrid screen that selected for compounds that prevent MTOR-Rictor protein-protein interaction and therefore specifically target mTORC2 signaling. This characteristic was further confirmed by multiple rounds of subsequent screening in the two-hybrid systems followed by analysis in mammalian cells. In addition, the small molecule inhibitors also lack any direct effect on a number of known kinases in biochemical assays (unpublished observations). In the Glioblastoma multiforme (GBM) tumor model, one of the isolated compound from the identical screen, CID613034 inhibited mTORC2 kinase activity in *vitro* and *in vivo* [27]. This compound inhibited the growth of GBM cell lines and the relative inhibition correlated with the expression of Rictor in the cells [27]. Immunoprecipitation experiments in also CLL specimen confirm the disruption of the mTORC2 complex as defined by the lack of co-immunoprecipitation of MTOR by Rictor antibody as a mechanism of action of these compounds.

Targeting BCR signaling and the activity of anti-apoptotic proteins has led to a number of newer therapeutic agents in this leukemia. A number of prior reports have shown that BCR signaling activates PI3Kinase/AKT signaling pathway and results in rapid phosphorylation of Akt. This Akt activation, in turn, upregulates a number of anti-apoptotic proteins such as Mcl-1, Bcl-xl, and XIAP [1–3,6–8]. The phosphorylation of Akt is observed in CLL specimens both with and without BCR crosslinking indicating that there is a baseline constitutive activation of this pathway in CLL specimens. The upregulation of pro-apoptotic proteins as mentioned above allows CLL specimens to resist apoptosis which is a hallmark of this leukemia. Blocking Akt phosphorylation by exposure to comp#6 results in altered signaling by Akt target genes namely GSK-3β, PKC-alpha, and FoxOs. Studies have shown that GSK3β phosphorylation and degradation can down-regulate the anti-apoptotic protein MCL-1 [7,36,37]. The pro-apoptotic Bim is also a target of GSK-3β as suppression of GSK-3β results in an increased expression of Bim in pancreatic cancer cells. Furthermore, GSK3β can phosphorylate and increase the activity and stability of BCL2L12a, an anti-apoptotic Bcl-2 family. PKC α is another downstream signaling target of Akt pathway and its signaling is also blocked by compound#6. Previous reports have shown a correlation between expression if its isoforms in CLL specimens and induction of apoptosis with PKC pathway inhibitors [38]. Additionally, another suggested mechanism of PKC-alpha is the phosphorylation of BCL2 in leukemia cells resulting in greater anti-apoptotic function [39].

Forkhead family of transcription factors (FoxOs) are phosphorylated and inactivated by Akt signaling and are known target genes of the PI3/Akt signaling pathway [40]. Blockade of Akt signaling results in an induction of apoptosis by either inducing expression of multiple pro-apoptotic members of the Bcl2-family or stimulating expression of death receptor ligands such as Fas ligand and tumor necrosis factor related apoptosis-inducing ligand (TRAIL) [41]. FoxOs induce Bim expression in hematopoietic cells deprived of growth factors and overexpression of a FoxO3a mutant that cannot be phosphorylated by Akt results in apoptosis of BCR-ABL-transformed cells [42]. The role of Forkhead family of transcription factors in the biology of CLL has not been reported so far. In this study we identified two forkhead members, FOXO1 and FOXO3A that are phosphorylated by BCR signaling. This phosphorylation is blocked by treatment with an mTORC2 inhibitor, compound#6, characterizing a role of forkhead family of transcription factors in protecting CLL cells against apoptosis

Stroma and microenvironment provides contact, cytokine and chemokine support to leukemic cells and resulting in resistance to apoptosis [8]. Interestingly both BTK inhibitor, Ibrutinib and PI3 kinase delta inhibitor result in initial lymphocytosis in CLL patients as they block microenvironment signaling and leukemic cells are no longer attracted to their niche in lymph nodes and bone marrow. Compound#6 was able to induce similar levels of apoptosis in CLL specimens both with and without stromal support implying that this compound has the ability to block microenvironment signaling. Similar activity of blocking microenvironment signaling is reported for the PI3 kinase inhibitor, Idelalisib as well [43].

Attempts have been made to block the activity of the PI3/Akt pathway at multiple levels to achieve growth inhibitory and apoptotic effects. Upstream inhibitors such as the PI3 kinase inhibitors Idelalisib, Duvesalib and Copanlisib are effective as their targets are over-expressed in B cell leukemia and lymphomas [44,45]. These inhibitors rely on the expression of specific isoforms of the PI3 kinase to be therapeutically active. Other classes of compounds which can inhibit mTORC2 activity include the ATP-competitive compounds such AZD8055 [46,47], mTORC1 inhibitors such as Temsirolimus and Everolimus, dual mTORC1/2 inhibitors and dual PI-3K/mTOR inhibitors, however, none of these inhibitors are very specific [19,20,48,49] and have the potential to block mTORC1 activity and result in the counter-productive activation of AKT. Compound#6 has a unique mechanism of action that specifically targets the mTORC2 complex signaling and with a greater ability to induce apoptosis in primary CLL leukemic specimens as compared to the PI3kinase inhibitor Idelalisib.

In summary, our findings indicate that the activity of the mTORC2 signaling complex is enhanced by Rictor over-expression in number of B cell malignancies. The compounds described in this report specifically and effectively abrogate mTORC2 signaling, an important B cell receptor-initiated signaling pathway by disrupting the mTORC2 complex. This is a novel mechanism which does not activate mTORC1 signaling and thereby does not allow paradoxical activation of Akt. In addition the inhibitors are distinct from PI3 kinase inhibitors as they do not rely on the expression of certain kinases. Further development of these compounds that block Rictor-mTOR interaction will be an effective therapeutic strategy for B cell malignancies.

## Acknowledgment

SS is supported by a grant from Flight Attendants Medical Research Institute (FAMRI) and Veterans Administration Merit Research award. JG has received funding from NIH R01CA109312 and Merit Award from Veteran Administration.

